# Systems-level Investigation of the Anxiolytic Gut–Brain Interactions induced by Paraprobiotic *Lactobacillus brevis* SBC8803 in Zebrafish

**DOI:** 10.64898/2026.02.04.703736

**Authors:** Azusa Kubota, Liqing Zang, Takuro Shinkai, Misa Nakai, Atsushi Tajima, Yasuhito Shimada

## Abstract

Anxiety disorders are among the most prevalent mental health conditions worldwide, and interest in psychobiotics—live or inactivated microorganisms that beneficially modulate the microbiota–gut–brain axis—is increasing. Heat-killed *Lactobacillus brevis* SBC8803 enhances serotonin (5-hydroxytryptamine; 5-HT) signaling and ameliorates stress-related phenotypes in mammals, although the gut–brain pathways mediating these effects remain incompletely defined. Here, we investigated the anxiolytic effects and underlying molecular mechanisms of oral SBC8803 administration in adult zebrafish.

Adult male AB-strain zebrafish were fed a diet containing heat-killed SBC8803 for 4 weeks, and anxiety-like behavior was evaluated using the novel tank test. SBC8803-treated fish exhibited a shorter latency to enter the upper half of the tank and more frequent transitions into the upper half, consistent with reduced anxiety-like behavior. To explore underlying mechanisms, we performed brain RNA sequencing and 16S rRNA gene sequencing of intestinal contents, followed by integrative multi-omics analyses. Brain transcriptomic profiling identified differentially expressed genes and enrichment of serotonin receptor, CREB, and oxytocin signaling pathways, suggesting enhanced monoaminergic and plasticity-related signaling. Microbiome functional prediction indicated SBC8803-associated shifts in lipid and vitamin metabolism, including pathways related to riboflavin (vitamin B2) and tryptophan.

Gene set variation analysis combined with DIABLO-based data integration revealed coordinated changes between microbial metabolic and brain signaling pathways, consistent with a vitamin B–serotonin–anti-inflammatory axis linking gut metabolism to neural regulation. Furthermore, residual correlation analysis showed innate gut–brain coordination independent of SBC8803 effect, such as the coupling between brain arachidonic acid and gut histidine metabolism. These findings support the biological validity of SBC8803 administration-associated interactions observed in the multi-omics analyses. Collectively, these findings indicate that the paraprobiotic SBC8803 exerts anxiolytic-like effects in zebrafish and reshapes gut–brain network states at behavioral, microbial, and transcriptomic levels, providing a mechanistic framework for considering heat-killed SBC8803 as a candidate psychobiotic for anxiety-related conditions.

## 1 Introduction

In 2019, an estimated 301 million people were living with anxiety disorders—approximately 4% of the global population—making them the most prevalent mental health conditions worldwide (WHO, *Anxiety disorders*. Final access on July 28, 2025. https://www.who.int/news-room/fact-sheets/detail/anxiety-disorders). The prevalence of anxiety disorders is expected to rise further. These disorders encompass generalized anxiety disorder, panic disorder, social anxiety disorder, specific phobias, selective mutism, separation anxiety, and agoraphobia (Ren et al., 2024). Affected individuals typically experience persistent or excessive anxiety, worry, and fear, often accompanied by avoidance of perceived threats and panic attacks, which substantially impair daily functioning and quality of life. First-line treatments include psychotherapy (Cuijpers et al., 2024; Sun et al., 2025) and pharmacotherapy (Bandelow et al., 2017; Kong and Han, 2024). Although artificial intelligence–based tools are being explored for anxiety management, concerns remain regarding privacy, algorithmic bias, and the need for human-centered approaches (Das and Gavade, 2024). Pharmacological options—such as selective serotonin reuptake inhibitors (SSRIs), serotonin–norepinephrine reuptake inhibitors (SNRIs), tricyclic antidepressants, and benzodiazepines—are effective (Luyten et al., 2011; Mohatt et al., 2014; Østergaard, 2018); however, limitations including delayed onset of action, adverse effects, and risk of dependency underscore the need for complementary strategies.

The gut–brain axis, a bidirectional network linking the central and enteric nervous systems with the immune system and the gut microbiota, plays a fundamental role in emotional and cognitive regulation (Aburto and Cryan, 2024; He et al., 2024). Dysbiosis has been associated with anxiety-related phenotypes, potentially through serotonergic, GABAergic, and immunomodulatory pathways (Foster et al., 2017; Jiang et al., 2024). Germ-free or antibiotic-treated animals exhibit exaggerated stress responses and anxiety-like behaviors, which can be mitigated by colonization with specific commensal microorganisms (Diaz Heijtz et al., 2011; Clarke et al., 2013). Preclinical and clinical studies further suggest that modulation of the gut microbiota can reduce anxiety symptoms. Randomized trials with probiotics have reported moderate improvements, particularly with *Bifidobacterium longum* and *Lactobacillus rhamnosus* (Mosquera et al., 2024), and multiple animal and human studies indicate reductions in stress- and anxiety-related measures following probiotic intake (Gholian et al., 2024). Collectively, these findings support the concept of psychobiotics—live or inactivated microorganisms that confer mental health benefits via the gut–brain axis—as promising adjunctive interventions.

Among candidate psychobiotics, *Lactobacillus brevis* SBC8803 is notable for its ability to enhance serotonin (5-hydroxytryptamine; 5-HT) signaling in the gut. This strain induces 5-HT release from enterochromaffin cells even in a heat-killed form (Nakaita et al., 2013). A non-randomized, double-blind, placebo-controlled crossover pilot study reported improved sleep quality following intake of heat-killed *L. brevis* SBC8803 (Nakakita et al., 2016). Additional studies have associated this strain to reduced stress-induced anxiety-like behavior and improved hippocampus-dependent memory (Higo-Yamamoto et al., 2019; Ishikawa et al., 2019).

The present study aimed to elucidate the molecular gut-brain pathways underlying the anxiolytic effects of heat-inactivated *L. brevis* SBC8803 using zebrafish as a translational model. Adult fish received oral SBC8803 and were assessed in the novel tank diving test. To investigate underlying mechanisms, we combined brain RNA sequencing with 16S rRNA gene profiling of the gut microbiota and conducted integrative multi-omics analyses to identify host transcriptomic changes and microbial functional alterations associated with anxiolytic-like behavior.

## 2 Materials and Methods

### 2.1 Ethics

All animal experiments were approved by the Ethics Committee of Mie University (approval number 2025-21) and were conducted in accordance with the Japan’s Act on Animal Welfare and Management of Animals (Ministry of the Environment of Japan). The experiments were also performed in compliance with the internationally accepted guidelines for the use of animals in research.

### 2.2 Zebrafish

Adult wild-type zebrafish (*Danio rerio*, AB strain, 6 months old; obtained from the Zebrafish International Resource Center, Eugene, OR, USA) were used in all experiments. Fish were maintained under standard laboratory conditions, including a 14:10 h light/dark cycle, a water temperature of 28.5 °C, and recirculating system with filtered tap water. Fish were group-housed (n = 10 per tank) in 2-L tanks and fed a commercial diet (Gemma Micro 300, Skretting, Stavanger, Norway) twice daily unless otherwise specified.

### 2.3 SBC8803 administration to zebrafish

To eliminate potential effects of sexual dimorphism, only adult male zebrafish were used in the feeding experiments. Five fish were housed in each 2 L tank. *Lactobacillus brevis* SBC8803 (FERM BP-10632) was obtained from the NITE Biological Resource Center (Kisarazu, Japan). SBC8803 was incorporated into Gemma Micro 300 powder at a concentration of 2.5 × 10^10^ cells per gram of feed. The mixture was lyophilized and subsequently processed according to a previously described method to achieve a particle size of ≤ 700 μm (Zang et al., 2011). Each fish received 5 mg of feed per feeding (equivalent to 1.25 × 10^8^ cells), administered twice daily for four weeks.

### 2.4 Novel tank diving test (NTT)

After the 4-week feeding period, anxiety-like behavior was assessed using the NTT. The NTT protocol consisted of three phases: acclimation, pre-test stress loading, and testing. During the acclimation phase, fish were placed in a 600-mL tank (Meito-Suien, Aichi, Japan; internal dimensions: 135 mm [width] × 95 mm [depth] × 90 mm [height]) for 5 min. Fish were then transferred to a holding tank for an additional 5 min to apply pre-test stress loading. Subsequently, fish were introduced into the test tank for behavioral assessment. The test tank had internal dimensions of 140 mm (width) × 45 mm (depth) × 120 mm (height). Behavioral recordings were captured at 30 frames per second (fps) for 5 min using a digital high-definition video camera (Panasonic HC-V495M, Osaka, Japan) positioned at a horizontal angle. Swimming behavior was analyzed using the iDTracker software (version 2.1) (Romero-Ferrero et al., 2019). According to the previous studies (Stewart et al., 2012; Shinkai et al., 2025), three parameters related to anxiety-like behavior were quantified: latency to enter the top half (LTTH), numbers of entries to the top half (NETH), and time spent in the top half (TSTH). In addition, total distance traveled and average swimming speed were measured.

### 2.5. Brain RNA-sequencing

Following behavioral testing, zebrafish were euthanized by ice water immersion, and whole brains were rapidly dissected and stored in RNAlater solution (Thermo Fisher Scientific, Waltham, MA, USA). Total RNA was extracted using TRIzol reagent (Thermo Fisher Scientific) in combination with the RNeasy Mini Kit for RNA cleanup (Qiagen, Hilden, Germany). RNA integrity was assessed using a Bioanalyzer 2100 system (Agilent Technologies, Santa Clara, CA, USA), and only samples with RNA integrity number (RIN) values greater than 8.0 were used for subsequent analyses. RNA libraries were prepared according to manufacturer’s protocol and sequenced using an MGI DNBSEQ-T7 platform.

### 2.6. RNA-seq data analysis

FASTQ files were trimmed by fastp (version 0.23.4) (Chen et al., 2018). Cleaned mRNA reads were aligned to the zebrafish reference genome using STAR (version 2.7.11b) (Dobin et al., 2013). The primary genome assembly FASTA file and corresponding genomic annotation GTF file of GRCz11 (Ensembl release version 113) was used for the alignment. Transcript abundance was quantified using Salmon (version 1.10.3) (Patro et al., 2017). For transcript quantification, the reference cDNA assembly was generated from the FASTA and GTF files using gffread (version 0.12.7) (Pertea and Pertea, 2020). Salmon output files were imported to R and converted into raw gene counts and normalized transcripts per million (TPM) values using tximport (version 1.34.0) (Love et al., 2018). TPM values were log_2_-transformed after adding a pseudocount 1 and used for principal component analysis (PCA). Differentially expressed gene (DEG) analysis was performed using DESeq2 (version 1.46.0) (Love et al., 2014). DEGs were identified using the following thresholds: false discovery rate (FDR) < 0.1 and absolute fold change ≥ 1.2. Annotation of DEGs was conducted using the R package biomaRt (version 2.62.1) (Smedley et al., 2009). Gene set enrichment analysis (GSEA) was performed using clusterProfiler (version 4.14.6) (Wu et al., 2021), with fold-change values derived from DESeq2 used as input. Gene sets with adjusted *p*-value < 0.1 were considered significant and categorized based on the sign of their normalized enrichment score (NES). Dot plots were generated to display up to the top 10 most statistically significant pathways for both the activated (NES > 0) and suppressed (NES < 0) gene sets. Molecular network and pathway analyses were further conducted using Ingenuity Pathway Analysis (IPA; QIAGEN, Redwood City, CA, USA). For IPA, genes with a nominal *p*-value < 0.01 in the DESeq2 results were used as input.

### 2.7 Intestinal 16S rRNA-sequencing

Intestinal contents were collected from the zebrafish after the behavioral testing, and gut bacterial DNA was extracted using a Quick-DNA Fecal/Soil Microbe Miniprep Kit (Zymo Research, Irvine, CA, USA). The V3–V4 region of the 16S rRNA gene was amplified by PCR, and library preparation and sequencing were performed using an Illumina MiSeq platform (2 × 300 bp). Sequence data were processed using QIIME2 (Hall and Beiko, 2018), and taxonomic classification was performed using the SILVA 138 database. Functional profiles of the microbiome were inferred using Phylogenetic Investigation of Communities by Reconstruction of Unobserved Status 2 (PICRUSt2) version 2.3.0 (Douglas et al., 2020).

### 2.8 Gene set variation analysis (GSVA)

Brain transcriptomic data and microbial PICRUSt2 output data were first converted into comparable pathway-level scores using GSVA implemented in the GSVA R package (version 2.0.7) (Hänzelmann et al., 2013). KEGG pathway gene sets for *Danio rerio* (mapped via Entrez Gene IDs) and microbial functions (mapped via KEGG Orthology identifiers) were retrieved using the KEGGREST R package (version 1.46.0). GSVA scores were calculated for each sample using the gsva function with a Gaussian kernel (kcdf = "Gaussian"), a minimum gene set size of 5, and a maximum size of 500. The resulting GSVA score matrices were used as the input for subsequent integration analyses.

### 2.9 Integrative multivariate analysis using DIABLO

Supervised multivariate analysis was performed using the DIABLO (Data Integration Analysis for Biomarker Discovery using Latent Variable Approaches for Omics Studies) framework implemented in the mixOmics R package (version 6.30.0) (Rohart et al., 2017). The GSVA score matrices for the brain (transcriptome block) and gut (microbiome_KO block) were used as predictor variables (X), with the experimental group (control vs SBC8803) specified as the outcome variable (Y). A design matrix specifying a between-block weight of 0.1 was applied, and the number of components (ncomp) was set to 1. The optimal number of pathways to retain from each block (keepX) was determined using the tune.block.splsda function, which performed 5-fold cross-validation repeated 10 times based on the Mahalanobis distance. The search grid for keepX ranged from 10 to 50 for the transcriptome block and from 5 to 25 for the microbiome_KO block. The final block.splsda model was constructed using the optimized keepX values (20 for the transcriptome block and 25 for the microbiome_KO). Pathway loadings on component 1, representing the contribution of each selected pathway to the discrimination, were subsequently extracted.

### 2.10 Analysis of residual correlations between brain and gut pathways

Residuals adjusted for the experimental group were first calculated for each pathway from the transposed GSVA score matrices (samples × pathways) for both transcriptome and microbiome_KO blocks using linear models (lm(pathway_score ∼ group_factor)). These residuals represent variation in pathway scores after accounting for the average effect associated with the experimental group, thereby isolating inter-individual variation independent of the main dietary intervention effect. Pairwise Spearman rank correlation coefficients and corresponding *p*-values were then computed between all brain and gut pathway residuals using the rcorr function from the Hmisc R package (version 5.2-3). The resulting *p*-value matrix was adjusted for multiple testing using the Benjamini-Hochberg procedure (p.adjust function, method = "fdr"), with an FDR < 0.05 considered statistically significant. To evaluate the robustness of these findings, a sensitivity analysis was performed in which the entire analysis pipeline, including GSVA and residual calculation, was repeated using five different pseudocount parameter values (1 × 10⁻⁷, 1 × 10⁻⁶, 1 × 10⁻⁵, 1 × 10⁻⁴, and 1 × 10⁻³) applied during the initial centered log-ratio (CLR) transformation of the KO abundance data (Gloor et al., 2017; Wang and Luan, 2024).

### 2.11 Statistical analysis

Behavioral and diversity data were analyzed using GraphPad Prism (version 10.5) and R (version 4.4.3). Statistical significance was determined using unpaired *t*-tests or one-way ANOVA, as appropriate. For RNA-seq and microbiome datasets, FDR-adjusted *p*-values < 0.05 were considered statistically significant. All data presented in graphs are expressed as mean ± standard deviation (SD).

## 3 Results

### 3.1 L. brevis SBC8803 shows anxiolytic effects in zebrafish NTT

After 4 weeks of SBC8803 administration, no mortality or significant changes in body weight were observed (**Figure S1**). In the NTT, SBC8803 significantly reduced latency to the top half (LTTH; *p* < 0.05; **Figure 1A**) and significantly increased numbers of entries to the top half (NETH; *p* < 0.01; **Figure 1B**). In contrast, no significant difference was detected in time spent in the top half (TSTH; **Figure 1C**). Under control conditions, zebrafish typically exhibit stress-induced dwelling in the lower half of the tank when exposed to a novel environment. SBC8803-administrated fish showed reduced anxiety-like responses, as indicated by their shorter LTTH and increased NETH. Although neither total distances traveled nor explored rate differed significantly between groups, both parameters showed a tendency to increase in SBC8803-treated fish (**Figure 1D** and **E**). Given the absence of significant changes in locomotor activity and the concurrent reduction in anxiety-related indices, these trends likely reflect enhanced exploratory behavior rather than general psychomotor activation. Collectively, these findings indicate that SBC8803 administration attenuates anxiety-like behavior in zebrafish without inducing nonspecific hyperlocomotion.

**Figure 1.**
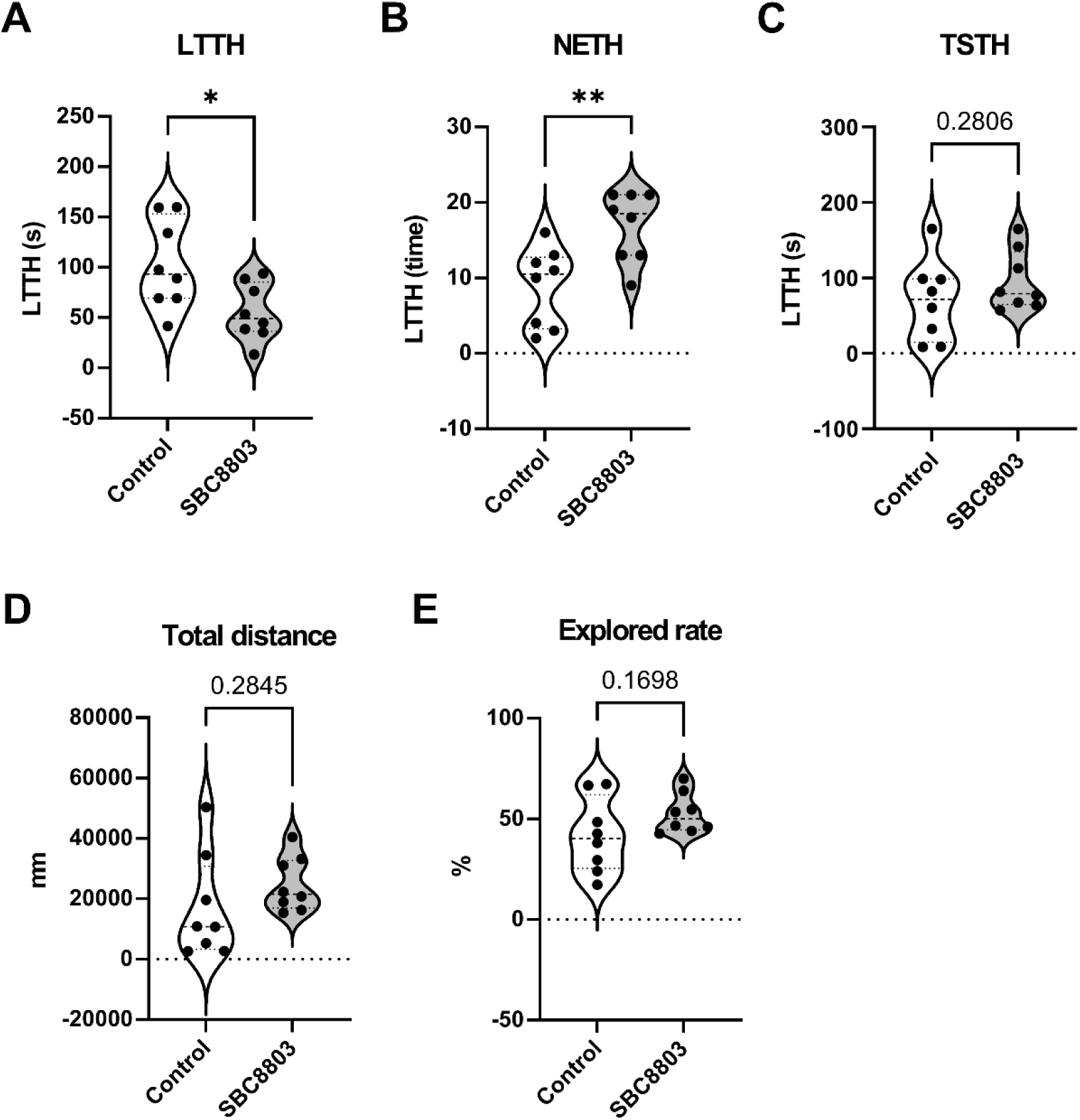
SBC8803 administration reduces anxiety-like behavior in the zebrafish novel tank test. (**A**) Latency to enter the top half of the tank (LTTH). SBC8803-treated fish exhibited a significantly shorter latency compared with control fish. (**B**) Number of entries into the top half of the tank (NETH). SBC8803 treatment significantly increased the frequency of transitions into the upper half. (**C**) Time spent in the top half of the tank (TSTH). No significant difference was observed between groups. (**D**) Total distance traveled during the test period. No significant difference was detected between control and SBC8803-treated fish. (**E**) Exploration rate during the novel tank test. Although SBC8803-treated fish showed a tendency toward increased exploration, the difference was not statistically significant. n = 8 per group. Data are presented as mean ± SD. \**p* < 0.05, \*\**p* < 0.01; ns indicate no significant difference.

### 3.2 Brain transcriptome of zebrafish treated with SBC8803

To investigate transcriptomic alterations associated with the anxiolytic phenotype, whole brains were collected following SBC8803 administration and subjected to RNA sequencing. For each sample, approximately 46–117 million paired-end reads were obtained, with high-quality scores across samples (Q20: 99.1–99.7%, Q30: 97.4–98.6%) and an average error rate of 0.02%, ensuring sufficient depth and accuracy for transcriptome analysis. PCA of log_2_-transformed TPM values was performed to visualize global variation between groups (**Figure S2**). Differential expression analysis using DESeq2 identified 128 DEGs, comprising 97 upregulated and 31 downregulated genes. These results are visualized in a volcano plot (**Figure S3)** and the complete list of DEGs is provided in **Table S1**. The full DESeq2 outputs is available in **Data S1**.

To interpret the biological relevance of these transcriptomic changes, we performed both over-representation analysis (ORA) and gene set enrichment analysis (GSEA) using DESeq2 results. For complementary pathway-level insight, a ranked gene list based on log_2_ fold change was subjected to GO-based GSEA. In total, 102 gene sets were enriched, of which 95 were activated and 7 were suppressed based on their normalized enrichment scores (NES; **Table S2** and **Figure 2A**).

**Figure 2.**
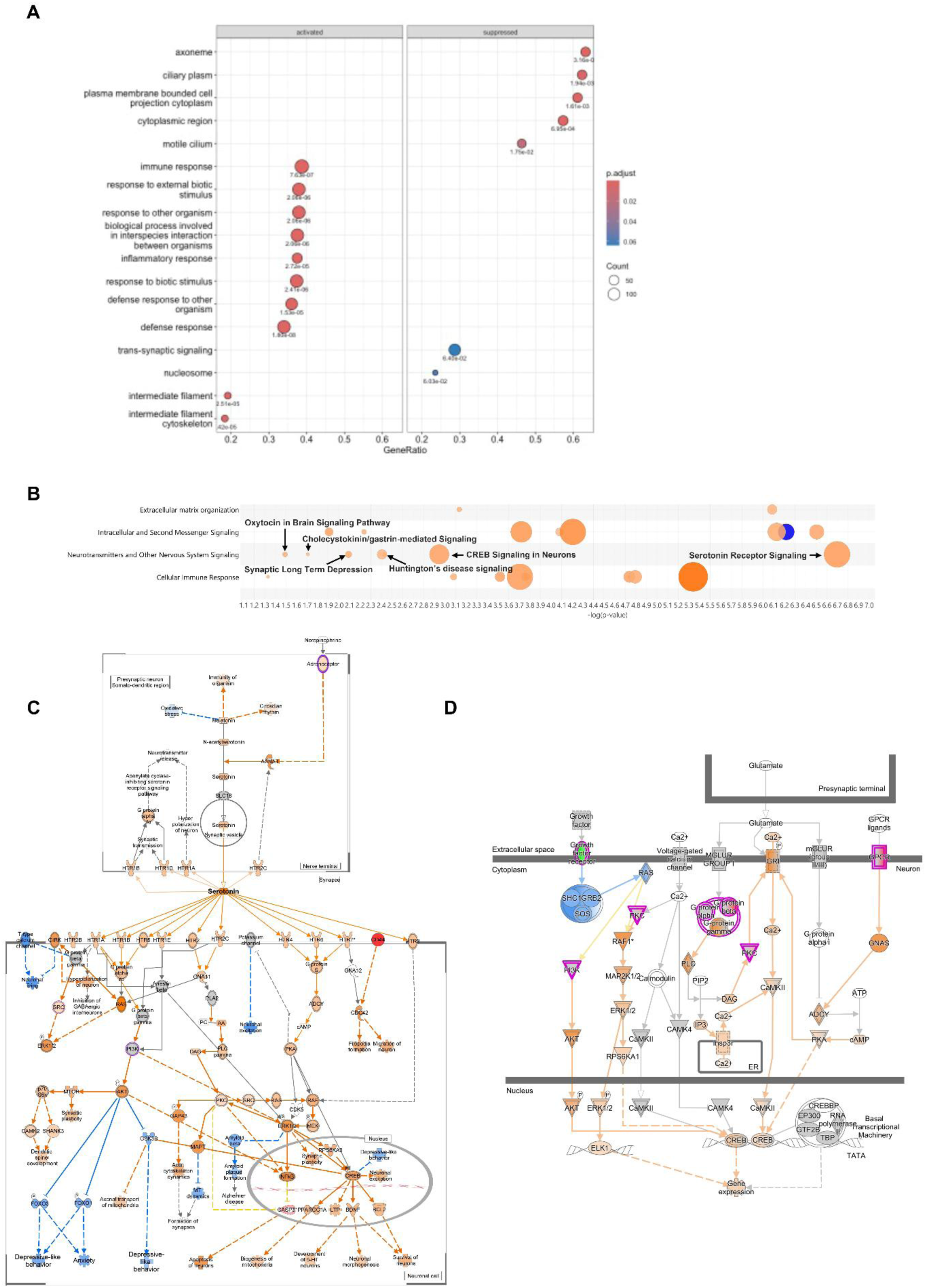
Brain transcriptomic alterations induced by SBC8803 administration. (**A**) Brain transcriptomic signatures identified by Gene Set Enrichment Analysis (GSEA). Gene Ontology (GO) enrichment analysis was performed using ranked log_2_(fold change) values derived from the DESeq2 analysis. The dot plots display the top significantly enriched GO terms (adjusted *p*-value < 0.1), categorized into activated (NES > 0) and suppressed (NES < 0) pathways. The x-axis represents the GeneRatio, defined as the proportion of genes contributing to the core enrichment subset relative to the total gene set size. The size of each dot corresponds to the number of enriched genes, and the color indicates the statistical significance (adjusted p-value). (**B**) IPA identifies enrichment of neurotransmitter-related signaling pathways, including serotonin receptor signaling, CREB signaling in neurons, and oxytocin signaling in the brain, which may underlie the anxiolytic-like effects observed following SBC8803 treatment. All enriched pathways are shown in **Figure S4**. (**C**) Predicted activation of the serotonin receptor signaling pathway in the zebrafish brain following SBC8803 administration. (**D**) Predicted activation of the canonical CREB signaling pathway in the zebrafish brain following SBC8803 administration. In panels C and D, genes with increased expression are shown in red, whereas genes with decreased expression are shown in green. Predicted activation and inhibition effects on signaling intermediates and downstream pathways are depicted by orange and blue lines, respectively, with darker hues indicating stronger predicted activation or inhibition.

We next applied Ingenuity Pathway Analysis (IPA) to identify signaling pathways modulated by SBC8803 administration (**Figure S4**). Using thresholds of –log(*p*-value) > 1.3 and z-score > 2, six pathways within the “Neurotransmitters and Other Nervous System Signaling” category were identified as activated. This included serotonin receptor signaling, CREB signaling in neurons, Huntington’s disease signaling, synaptic long-term depression, cholecystokinin/gastrin-mediated signaling, and oxytocin signaling in the brain (**Figure 2B**). Ranked pathways potentially associated with mental disorders were extracted based on IPA analysis following SBC8803 treatment, using a threshold of –log(*p*-value) > 1.3 (**Table 1**). Among these 14 canonical pathways, serotonin receptor signaling emerged as the most strongly activated pathway (–log(*p*-value) = 6.70, z-score = 2.828; **Figure 2C**). Key DEGs contributing to serotonin receptor signaling—including *GNA15*, *GNB3*, *ADRB3*, and *PRKCH*—were significantly upregulated in the DESeq2 analysis (fold change > 1.5, nominal *p*-value < 0.05) and were subsequently mapped to this pathway by IPA. These genes are involved in G protein-coupled receptor (GPCR) signaling and downstream second-messenger cascades associated with 5-HT1A/1B and 5-HT2 receptor function, which are well-established regulators of anxiety-related behaviors in both zebrafish and mammals (Connors et al., 2014; Wong et al., 2023).

Consistent with serotonergic enhancement, IPA also indicated activation of the cAMP response element-binding protein (CREB) signaling pathway (**Figure 2D**). Upstream regulators, including CaMKII, PKA, and ERK1/2, were either upregulated or predicted to be activated, converging on CREB phosphorylation and transcriptional activation. These changes are consistent with enhanced relevant to synaptic plasticity and transcriptional programs involved in emotional regulation.

### 3.3 Microbiota alteration by SBC8803 administration in zebrafish

To investigate the effects of SBC8803 on the intestinal microbiota, we profiled the gut microbiomes of control and SBC8803-treated zebrafish by amplicon sequencing of the V3–V4 region of the 16S rRNA gene. For each sample, 60,000–150,000 paired-end reads were obtained, with Q20 and Q30 scores of 98.0–98.5% and 93.4–94.9%, respectively. After quality filtering and processing with QIIME2, a total of 450 operational taxonomic units (OTUs) were identified. Despite continuous SBC8803 administration and its clear anxiolytic effect, alpha diversity was comparable between the two groups; Shannon diversity, Chao1 richness, and Pielou evenness indices showed no significant differences (**Figure 3A**). At the phylum level, the gut microbiota of both groups was dominated by Fusobacteria and Proteobacteria, in agreement with previous reports for adult zebrafish. Family-level profiles revealed a modest increase in *Fusobacteriaceae* and a marked decrease in Enterobacteriaceae in the SBC8803 group relative to controls (**Figure 3B**). At the genus level, no taxa displayed statistically significant differences after SBC8803 treatment (all *p* > 0.05), although *Levilactobacillus*, *Chitinilyticum*, and Flavobacterium tended to increase and *Plesiomonas* tended to decrease (**Figure 3C)**. Using PICRUSt2 for functional prediction, we found that SBC8803 administration significantly increased the MetaCyc pathway PWY-5971 (palmitate biosynthesis II) and significantly decreased PWY-6629 (superpathway of L-tryptophan biosynthesis) (**Figure 3D**). The top 10 taxa at each taxonomic level (phylum, class, order, family, genus, and species) are summarized in **Data S2**.

**Figure 3.**
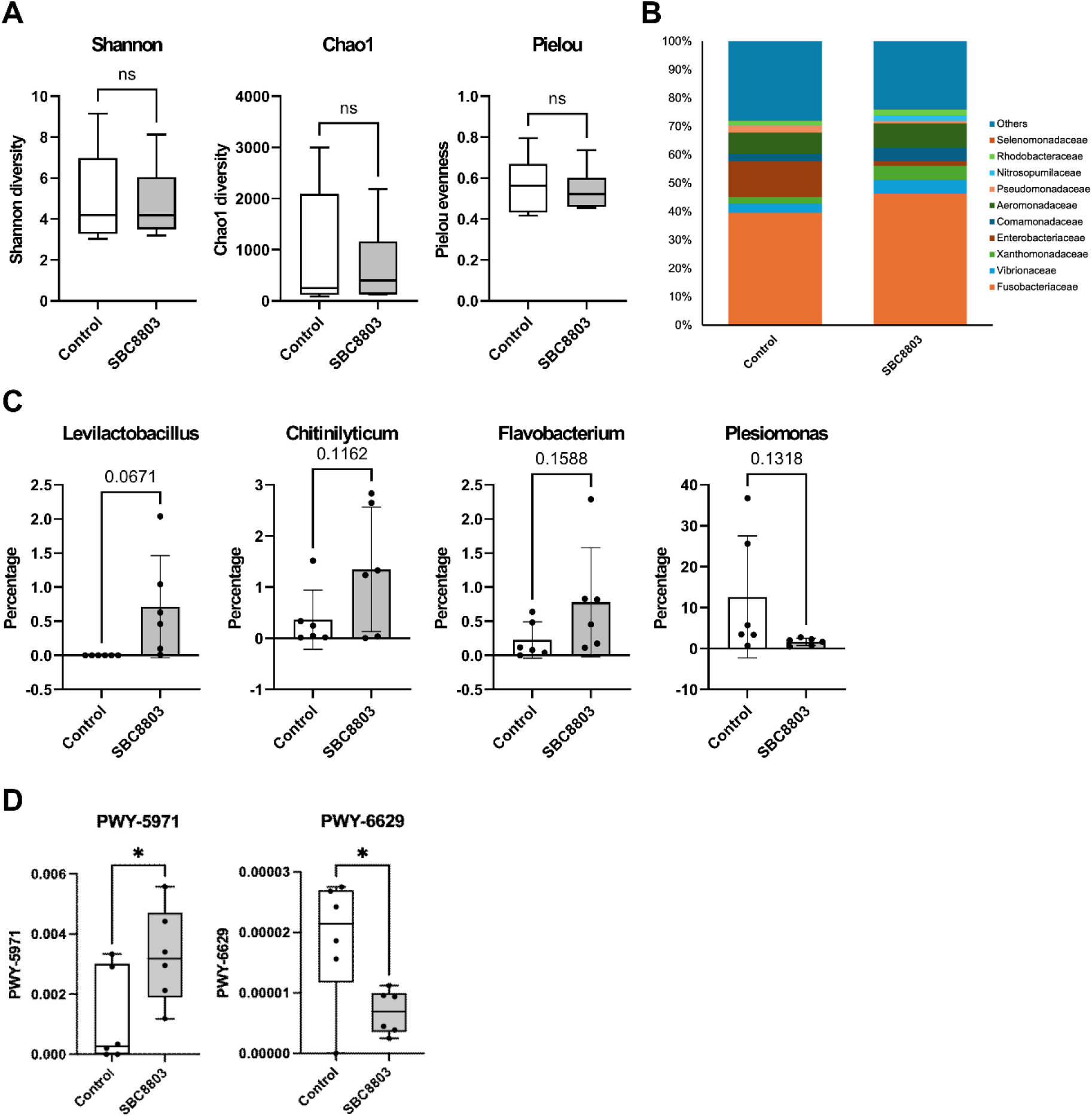
Effects of SBC8803 administration on the intestinal microbiota of zebrafish. (**A**) Alpha diversity of the gut microbiota in control and SBC8803-treated zebrafish, assessed using the Shannon diversity index, Chao1 richness, and Pielou evenness. No significant differences were observed between the two groups. (**B**) Relative abundance of dominant bacterial families in the gut microbiota of control and SBC8803-treated zebrafish. SBC8803 administration was associated with a modest increase in *Fusobacteriaceae* and a marked decrease in Enterobacteriaceae. (**C**) Relative abundance of bacterial genera in the gut microbiota of control and SBC8803-treated zebrafish. No genera exhibited statistically significant differences between groups (all *p* > 0.05), although *Levilactobacillus*, *Chitinilyticum*, and *Flavobacterium* tended to increase, whereas *Plesiomonas* tended to decrease following SBC8803 administration. (**D**) Functional prediction of microbial metabolic pathways inferred by PICRUSt2 and annotated using the MetaCyc database. SBC8803 administration significantly increased PWY-5971 (palmitate biosynthesis II) and significantly decreased PWY-6629 (superpathway of L-tryptophan biosynthesis).

### 3.4 Multi-omics integration reveals coordinated gut–brain metabolic signatures

Supervised integration of brain transcriptome and gut microbiome pathway scores DIABLO revealed a latent component structure strongly associated with the experimental groups (**Figure 4A**). The performance and robustness of the DIABLO model were assessed using M-fold cross-validation (5 folds, 10 repeats) during the tuning procedure. This analysis identified an optimal feature set consisting of 20 pathways from the transcriptome block and 25 pathways from the microbiome_KO block (keepX = 20 and 25, respectively), yielding a minimum mean classification error rate of 2.5% (**Figure S5**).

**Figure 4.**
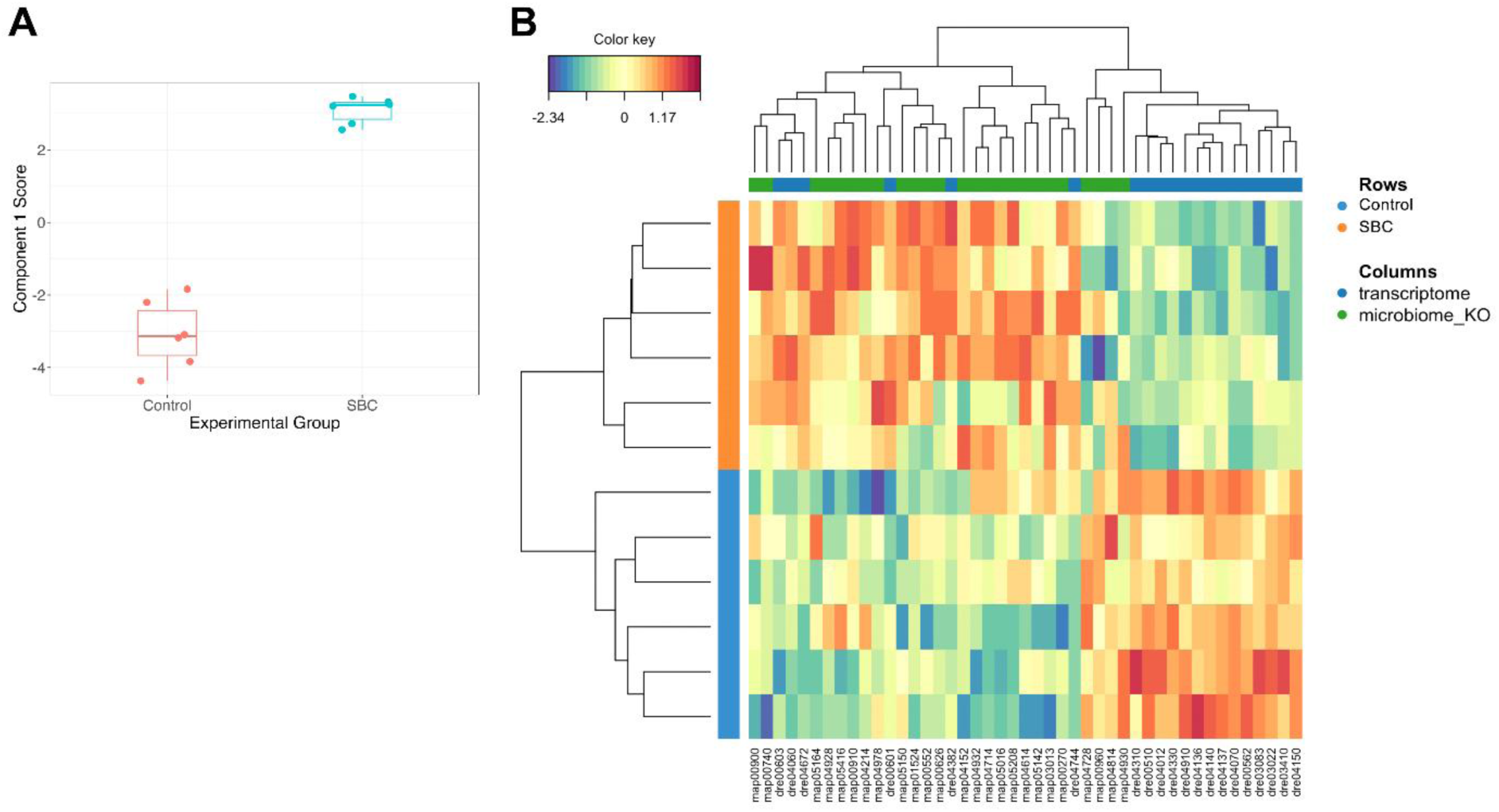
Multi-omics discrimination and molecular profiling using DIABLO. (**A**) Sample score plot for Component 1. The latent component scores, derived from the DIABLO (N-block sPLS-DA) framework, summarize the maximal covariance between the gut microbiome and brain transcriptome datasets. (**B**) Heatmap representing the cross-omics molecular profiles. The heatmap displays standardized GSVA scores for the top-ranking discriminative features selected by the model, revealing integrative classification and hierarchical clustering or the experimental groups

To visualize the cross-omics integration, correlations among the selected features from both data blocks were displayed as a circos plot (**Figure 5A**). Among the brain pathways, mTOR signaling pathway (dre04150; loading: −0.486) contributed most strongly to the control group, whereas cytokine-cytokine receptor interaction (dre04060; loading: +0.335) was most strongly associated with the SBC8803-treated group (**Figure 5B**). In the gut microbiome, the SBC8803 group was characterized by increased contributions from cysteine and methionine metabolism (map00270; loading: +0.420) and riboflavin metabolism (map00740; loading: +0.306) (**Figure 5C**).

**Figure 5.**
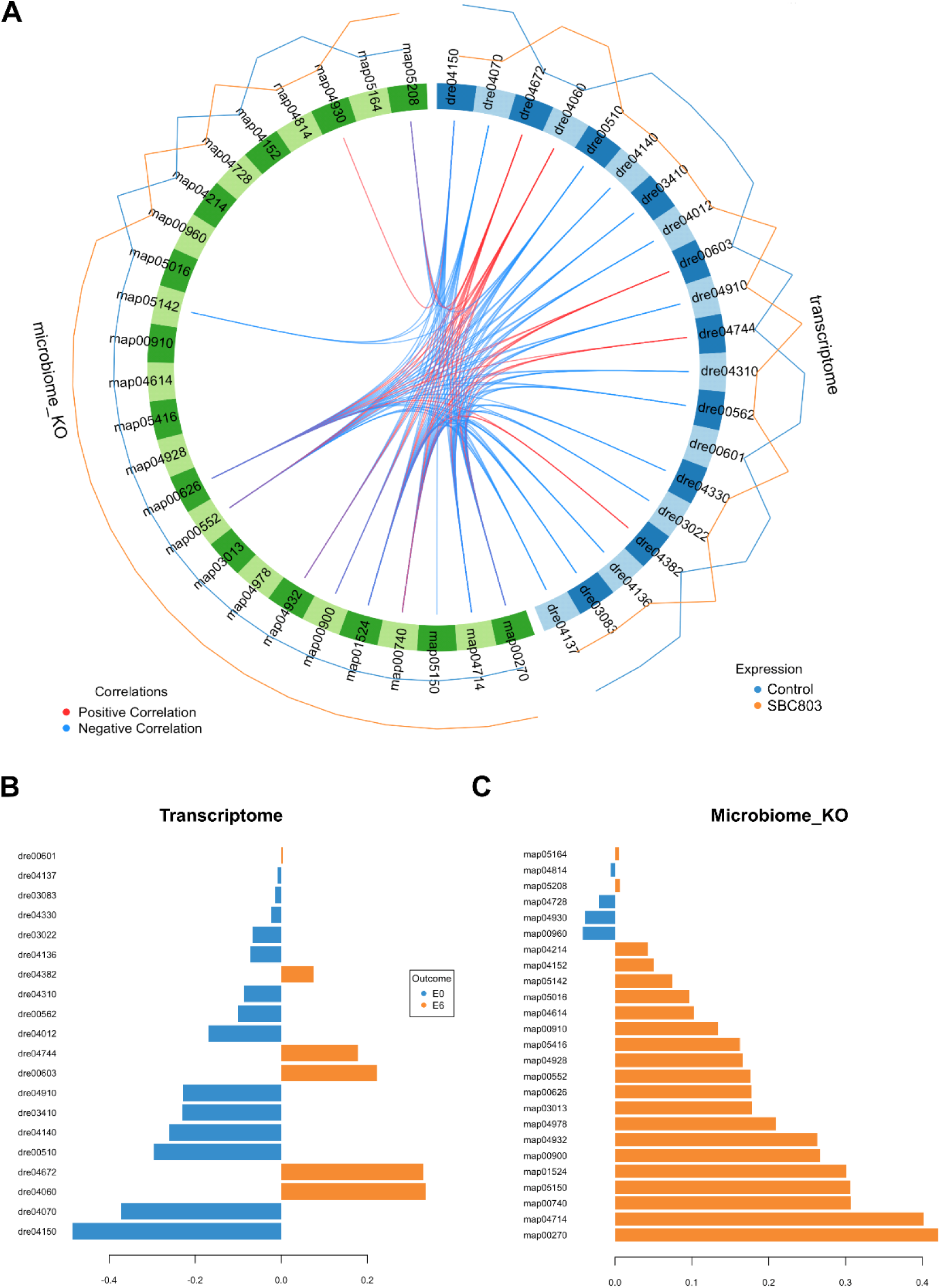
Discriminative pathway signatures and inter-modular cross-omics correlations. Identification of key microbial and brain transcriptomic features contributing to group separation and their functional associations. (**A**) Circos plot illustrating correlations between the most discriminative brain and gut pathways. Links represent strong Spearman correlations (|ρ| > 0.7), highlighting coordinated shifts between host neural pathways and microbial functional potentials identified by the DIABLO model. (**B–C**) Loading plots for the brain transcriptome (**B**) and gut microbiome (**C)**. In both omics blocks, KEGG Pathway identifiers are ranked according to their loading weights for Component 1, which represents the primary axis of separation between the control and SBC8803 groups. Colors indicate the group in which pathway activity is highest.

### 3.5 Observation of intrinsic gut–brain coordination independent of treatment effects

To confirm the association between the gut–brain interactions observed by DIABLO analysis and fundamental physiological coupling, we complementarily performed Spearman correlation analysis using residuals obtained after regressing out the control/SBC8803 group effect (**Figure 6A**). This approach isolates inter-individual variation independent of the dietary intervention, allowing us to detect robust pathway pairs that covary across the population regardless of the treatment status. A global heatmap summarizing all associations (**Figure S6**), and total 385 significant gut–brain pathway pairs were identified (FDR < 0.05; **Data S3**).

**Figure 6.**
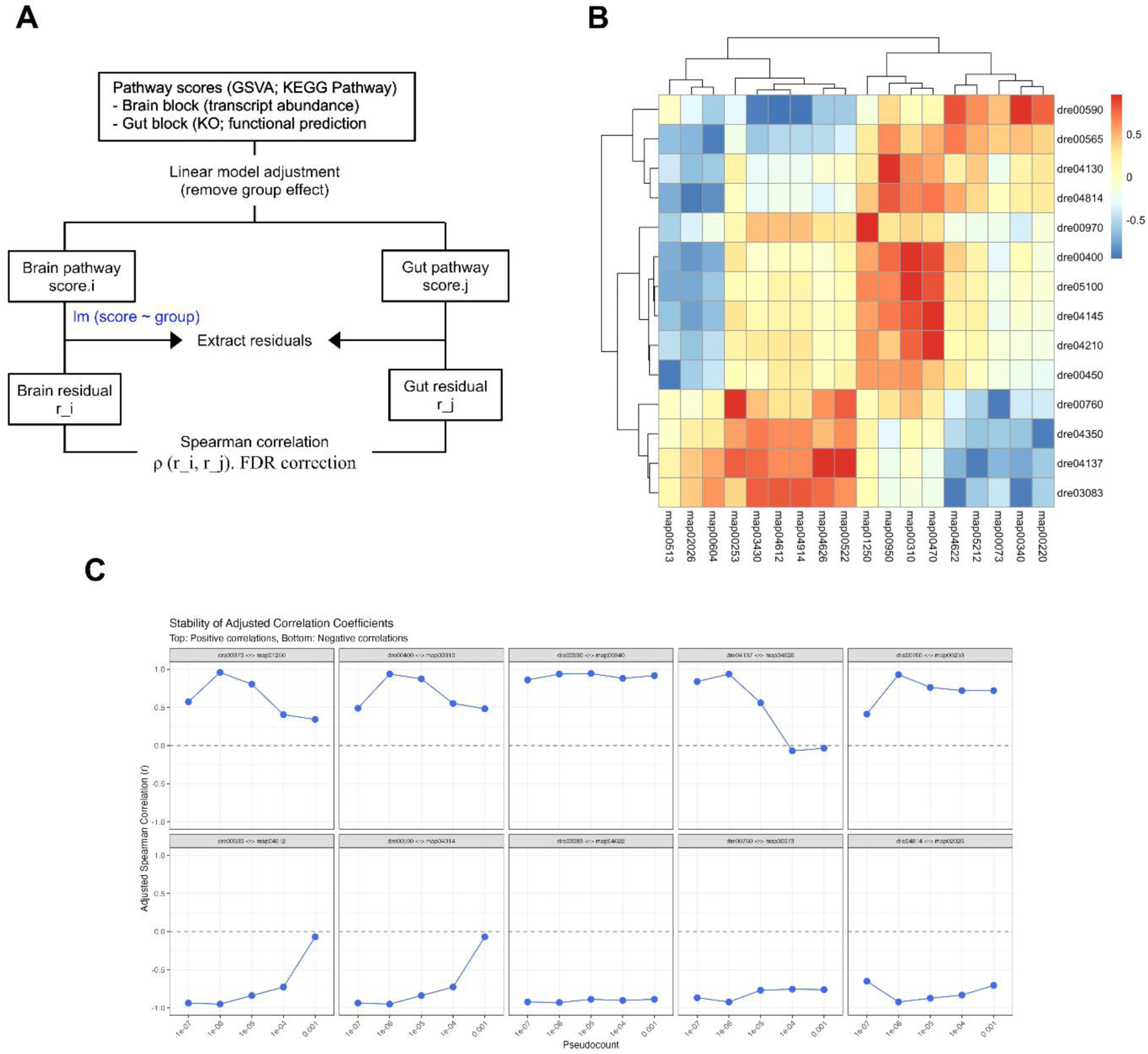
Gut–brain coordination independent of group effects shown by residual-based correlation analysis. (**A**) Conceptual overview of the residual correlation analysis workflow. To isolate gut–brain associations independent of the dietary intervention, pathway scores were adjusted using a linear model to regress out the group effect, and the resulting residuals were used for correlation analysis. (**B**) Heatmap of significant residual correlations between gut and brain pathways. Spearman’s correlation coefficients were calculated using residuals obtained after regressing out the group effect. Positive (red) and negative (blue) values indicate direct and inverse associations, respectively, independent of group-driven variations. (**C**) Sensitivity analysis of correlation stability. Plots demonstrate the robustness of adjusted Spearman’s correlation coefficients across varying pseudocount thresholds applied during the centered log-ratio (CLR) transformation.

To focus on the most robust associations, we examined the top ten positive and negative correlations ranked by effect size (**Figure 6B**). Among the positive associations, brain arachidonic acid metabolism (dre00590) and gut histidine metabolism (map00340) exhibited a strong correlation (ρ = +0.94, FDR = 0.0040) together with high stability across pseudocount settings (correlation range ≈ 0.08; **Figure 6C**). Among the negative associations, the most stable pair linked brain polycomb repressive complex–related activity (dre03083) with gut RIG-I-like receptor signaling (map04622), showing a strong inverse correlation (ρ = −0.93, FDR = 0.0053) with minimal variation in the sensitivity analysis (correlation range ≈ 0.04). Other pathway pairs, including the association between brain phenylalanine/tyrosine/tryptophan biosynthesis (dre00970) and gut secondary metabolite biosynthesis (map01250), displayed even higher coefficients (ρ = +0.96, FDR = 0.00084) but showed lower stability across pseudocount conditions (correlation range ≈ 0.62) and were therefore interpreted with greater caution.

## 4. Discussion

This study provides an integrative view of how heat-killed *L. brevis* SBC8803 can exert psychobiotic effects through multi-layered interactions within the microbiota–gut–brain axis. Multi-omics integration revealed coordinated alterations in gut microbial function and brain transcriptional regulation, indicating a systems-level network linking microbial vitamin metabolism with host neurotransmission and immune balance. Residual correlation analysis subsequently indicated potential gut–brain association independent of SBC8803 administration, supporting the interpretation of the multi-omics analyses. Together, these findings propose a model of a metabolically mediated gut–brain crosstalk in which microbial activity related to vitamin B metabolism converges with serotonergic signaling to shape neural and immune homeostasis.

PICRUSt2-based functional prediction illuminated the initial metabolic shifts in the gut. Palmitate biosynthesis (map00062: palmitate biosynthesis II) was upregulated, whereas the L-tryptophan biosynthesis superfamily was suppressed in the *L. brevis* SBC8803-supplemented group. Elevated microbial palmitate biosynthesis suggests a shift toward fatty acid anabolism and membrane lipid remodeling that may influence host immune tone through altered microbial metabolite profiles. Because fatty acid synthesis requires substantial NADPH and flavin cofactors (Byrnes et al., 2018; Tolomeo et al., 2020), this pattern is consistent with the observed activation of riboflavin (vitamin B2) metabolism and redox regulation. In contrast, suppression of microbial tryptophan biosynthesis implies reduced de novo capacity, plausibly reflecting increased host utilization of tryptophan for the serotonin and kynurenine pathways. This complementary relationship between microbial lipid metabolism and host neurotransmitter synthesis indicates a coordinated redistribution of metabolic resources along the microbiota–gut–brain axis, aligning with the proposed “vitamin B–serotonin–anti-inflammatory” model.

DIABLO multivariate analysis revealed a strong negative association between gut-derived riboflavin metabolism and brain mTOR signaling. In parallel, IPA identified activation of serotonin receptor and CREB signaling pathways in the supplemented group, suggesting enhanced synaptic plasticity-related signaling. Combined with GSEA-identified suppression of trans-synaptic signaling, these changes indicate a shift toward neural adaptation under the supplementation. This pattern is consistent with the idea that increased microbial vitamin B metabolism, particularly riboflavin, can indirectly modulate brain mTOR activity while supporting neurotransmission via serotonergic and CREB pathways (Udhayabanu et al., 2017; Ilchibaeva et al., 2022; Olfat et al., 2022). It also aligns with previous findings that vitamins B2 and B6 support redox balance and mitochondrial homeostasis, thereby mitigating stress-induced mTOR hyperactivation (Johnson et al., 2013; Ciapaite et al., 2023). Notably, IPA highlighted upregulation of serotonin receptor signaling, and DEG analysis detected increased expression of *tcnbb* (transcobalamin beta b), consistent with a role of vitamin B-dependent tryptophan hydroxylase in serotonin synthesis (Kennedy, 2016; Young et al., 2019; Tang et al., 2025). Collectively, these results point to a metabolically coordinated interaction between gut-derived vitamins and host neural signaling pathways, highlighting a mechanism by which microbial metabolism shapes brain homeostasis in zebrafish.

Beyond metabolic correlations, significant upregulation of *nfkbie*, an inhibitor of NF-κB, supports activation of an anti-inflammatory regulatory program in the brain. The inferred inhibition of NF-κB-related pro-inflammatory genes suggests that paraprobiotic treatment engages tolerance-like mechanisms that mitigate neuroinflammation. This interpretation is consistent with the GO GSEA results, where enrichment of the broad ‘inflammatory response’ and ‘immune response’ gene sets likely reflects regulatory engagement by anti-inflammatory mediators rather than a purely pro-inflammatory cascade. Concomitant suppression of mTOR and MAPK signaling further reinforces this interpretation, as both pathways mediate inflammatory amplification and neuronal excitotoxicity (Switon et al., 2017; Canovas and Nebreda, 2021). Taken together, these data support a model in which microbial vitamin B metabolism indirectly enhances serotonergic activity and concurrently dampens inflammation, stabilizing neural circuits.

Previous studies provide mechanistic context for this framework. The “vitamin B–serotonin–anti-inflammatory axis” is increasingly recognized as a conduit in microbiota–gut–brain communication. Vitamins B6 and B9 (folate) act as cofactors in the enzymatic conversion of tryptophan to serotonin in the gut epithelium (Wan et al., 2022; Lu et al., 2024). Approximately 90% of total serotonin is synthesized in the gastrointestinal tract, where microbial and host metabolism jointly regulate tryptophan hydroxylase expression (González Delgado et al., 2022; Miri et al., 2023). Gut-derived serotonin not only modulates intestinal motility but also signals via vagal afferents to the brain, influencing affective and cognitive processes (Horn et al., 2022). On immune cells, activation of 5-HT2B and 5-HT4 receptors promotes an anti-inflammatory phenotype (Wan et al., 2020; Zheng and Xu, 2025); in the brain, 5-HT1A and 5-HT2A receptors activation suppresses microglial activation and oxidative stress (Krabbe et al., 2012; Glebov et al., 2015; Xing et al., 2024). Therefore, the observed upregulation of serotonergic signaling and the concurrent activation of an anti-inflammatory response (evidenced by *nfkbie* upregulation) in our zebrafish model may underlie behavioral stabilization via suppression of neuroinflammatory tone (Whiteside et al., 1997; Demin et al., 2021). These data collectively indicate that the vitamin B–serotonin–anti-inflammatory axis not only promotes molecular homeostasis but also contributes to behavioral resilience mediated by the gut–brain interaction.

Residual correlation analysis, designed to observe the gut–brain associations independent of SBC8803’s effects, revealed two notable patterns. The first was a stable negative correlation between the brain polycomb repressive complex pathway (dre03083) and the gut RIG-I-like receptor signaling pathway (map04622). The gut RIG-I signal here represents a predicted microbial functional feature (homologs of dsRNA-recognition and RNA helicase genes) rather than activation of the host pathway. In light of the GSEA-indicated suppression of the nucleosome gene set, suggesting a shift toward less compact chromatin, reduced polycomb activity is plausible. If so, the inverse correlation implies enhanced bacterial antiviral-like functional potential in the supplemented group alongside reduced PRC activity in the brain. Given PRC1/PRC2 roles in neural development and glial maintenance in zebrafish (Raby et al., 2021; Feng and Sun, 2022; Hanot et al., 2025), this coordinated pattern may reflect a compensatory mechanism whereby microbial immune-like functional states interface with central epigenetic regulation (Liu et al., 2023). The second notable finding was a robust positive association between brain arachidonic acid metabolism (dre00590) and gut histidine metabolism (map00340). Given that both arachidonic acid–derived lipids and histidine-derived metabolites modulate neuroimmune signaling (Broos et al., 2024; Dürholz et al., 2025), this covariation suggests an additional axis of coordinated gut–brain functional variation that persists beyond group-level differences. Taken together, these two findings are consistent with biological features not captured by DIABLO-based multi-omics integration, supporting the view that the paraprobiotic SBC8803 may exert additional effects beyond canonical gut–brain coupling.

Our results also suggest that paraprobiotics can elicit systemic effects comparable to those of live probiotics, in line with recent evidence. Multiple studies report that heat-killed *Lactobacillus* strains (e.g., *L. mucosae*, *L. plantarum*, *L. rhamnosus*) increase colonic IL-10, modulate serotonin receptor expression, enhance 5-HT biosynthesis, and suppress pro-inflammatory cytokines (Magryś and Pawlik, 2023; Mudaliar et al., 2024; Sun et al., 2024). These data support the view that nonviable microbial components can engage host pattern-recognition receptors and metabolic circuits, producing beneficial psychobiotic effects via immunometabolic modulation.

Several limitations should be noted. First, the associations reported here are correlational and do not establish causality; targeted metabolite supplementation or microbial genetic manipulations will be required to test causality. Second, microbial functions were predicted from 16S rRNA gene data and may not fully reflect activity; metabolomic measurements of intestinal contents and serum are needed to confirm production and circulation of vitamin-B derivatives and serotonin precursors. Third, while zebrafish provide valuable translational insight, extrapolation warrants caution; studies in rodents or humanized models will be necessary to evaluate the generalizability of the proposed vitamin B axis and its behavioral consequences.

In summary, our data support a mechanistic framework in which gut-derived vitamins, serotonergic signaling, and anti-inflammatory pathways cooperatively sustain neural and behavioral stability. This integrated view reinforces the concept of paraprobiotics as effective psychobiotic modulators and provides a foundation for future therapeutic strategies targeting the microbiota–gut–brain axis.

## 5. Conclusion

This study demonstrates that paraprobiotic supplementation reshapes the gut–brain network through the coordinate activation of microbial vitamin B metabolism and serotonergic pathways, leading to the suppression of neuroinflammatory signaling. The integrative multi-omics approach highlights a systemic “vitamin B–serotonin–anti-inflammatory axis” linking gut metabolic function to neural gene regulation, providing a novel mechanistic basis for the psychobiotic potential of probiotics.

## Supporting information

The complete list of differentially expressed genes (DEGs)

The full DESeq2 outputs

The top 10 taxa at each taxonomic level (phylum, class, order, family, genus, and species)

List of enriched Gene Ontology (GO) terms identified by Gene Set Enrichment Analysis (GSEA)

## Table

**Table 1.** Canonical pathways associated with mental disorders identified by IPA

## Supplementary Figures

**Figure S1.** Effects of SBC8803 administration on somatic growth parameters in adult zebrafish.

**Figure S2.** Principal component analysis (PCA) of brain transcriptomes.

**Figure S3.** Volcano plot of differentially expressed genes (DEGs) between the control and SBC8803-treated groups.

**Figure S4.** Canonical signaling pathways modulated by SBC8803 administration.

**Figure S5.** Model evaluation and parameter tuning for the DIABLO classification.

**Figure S6.** Overall adjusted correlation between brain transcriptome and gut microbiome pathways.

**Figure S7.** Scatter plots of significant residual correlations between gut microbial and brain pathways.

## Supplementary Tables

**Table S1.** The complete list of differentially expressed genes (DEGs)

**Table S2.** List of enriched Gene Ontology (GO) terms identified by Gene Set Enrichment Analysis (GSEA)

## Supplementary Data

**Data S1.** The full DESeq2 outputs.

**Data S2.** The top 10 taxa at each taxonomic level (phylum, class, order, family, genus, and species).

**Data S3.** Significant gut–brain pathway pairs identified by residual correlation analysis.

## Acknowledgements

We thank Ms. Masami Yamamura for her technical assistance with zebrafish breeding and husbandry, and Ms. Rie Ikeyama for her secretarial support.

## Funding

This study was supported by JST SPRING (grant number JPMJSP2135).

## Conflicts of interest

Y.S. is the CTO and an executive member of Zebra-Innovate LLC. The other authors declare that the research was conducted in the absence of any commercial or financial relationships that could be construed as a potential conflict of interest.

## Data Availability Statement

The datasets presented in this study can be found in online repositories. The names of the repository/repositories and accession number(s) can be found below: NCBI Gene Expression Omnibus (GEO), accession numbers GSE314984 (Brain RNA-seq) and GSE314985 (Intestinal 16S rRNA-seq).

