## Supplementary Figures for "Systems-level Investigation of the Anxiolytic Gut–Brain Interactions induced by Paraprobiotic *Lactobacillus brevis* SBC8803 in Zebrafish"

**Figure S1**

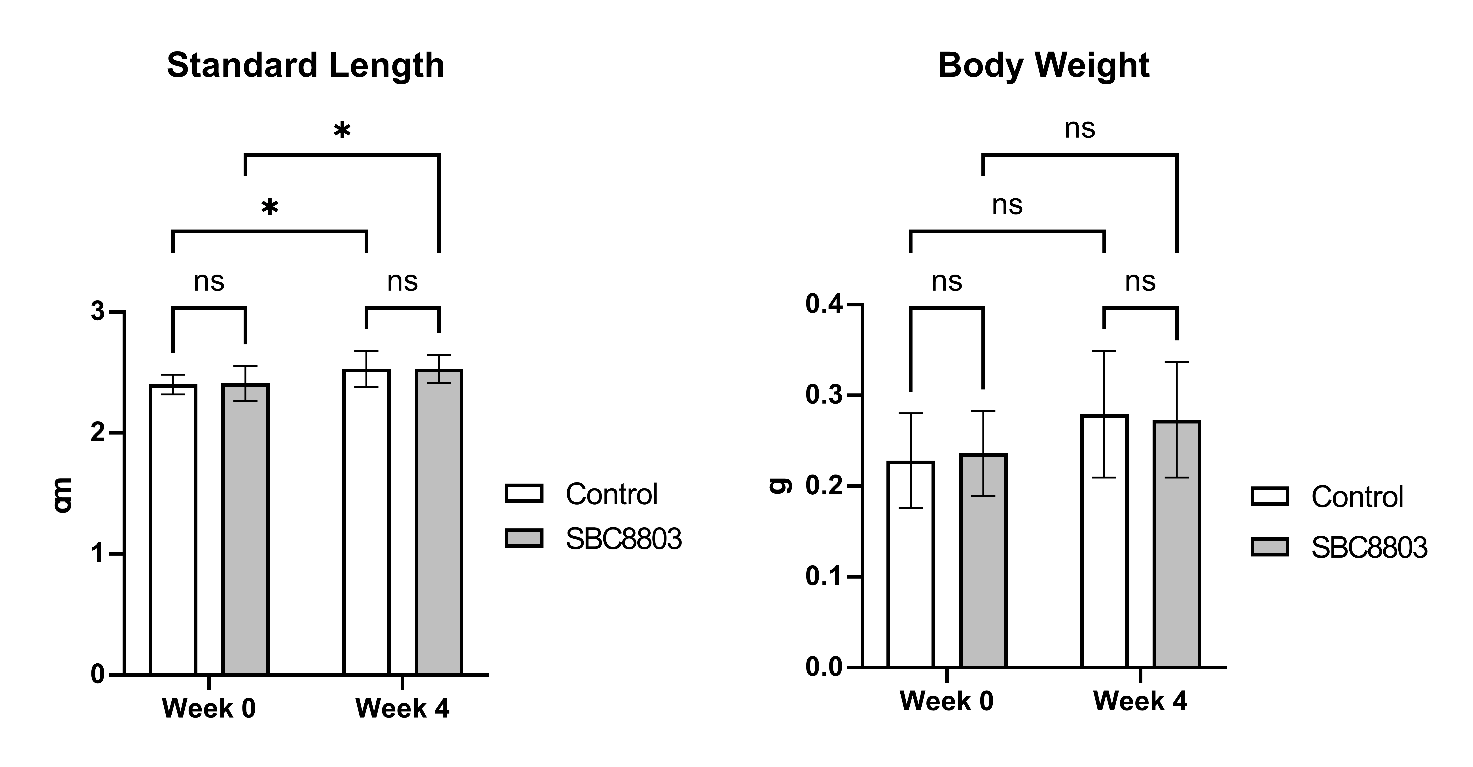

**Figure S1. Effects of SBC8803 administration on somatic growth parameters in adult zebrafish.** Standard length (left) and body weight (right) were measured before (week 0) and after (week 4) the 4-week feeding period. Both the control and SBC8803-treated groups exhibited a significant increase in standard length over time (* *p* < 0.05 vs. Week 0), reflecting normal growth. No significant differences were observed between the control and SBC8803 groups in either standard length or body weight at any time point, indicating that SBC8803 administration did not negatively affect physical development. Data are presented as mean ± SD (n = 8 per group). ns, not significant; * *p* < 0.05.

**Figure S2**

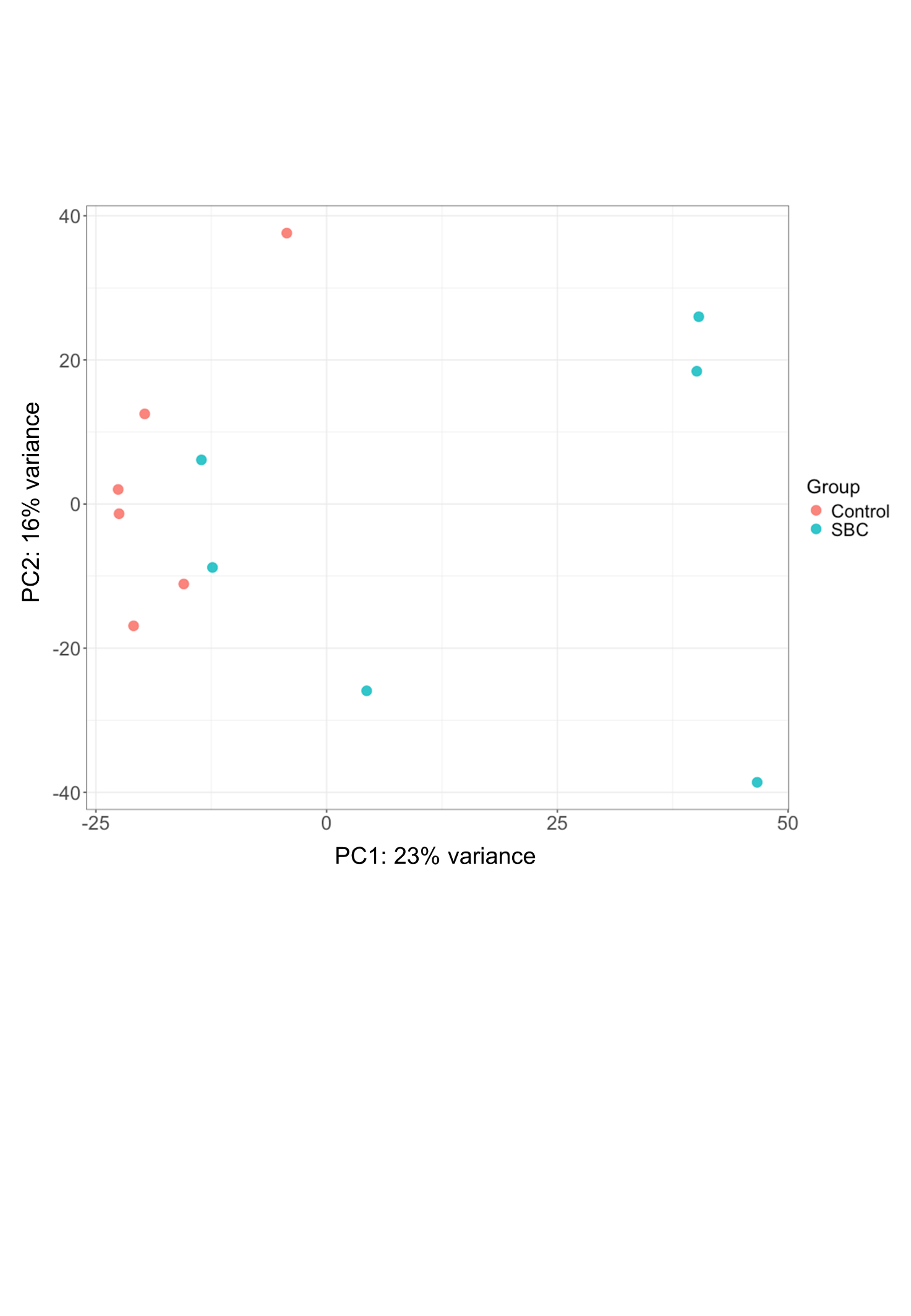

**Figure S2. Principal component analysis (PCA) of brain transcriptomes.** A scatter plot visualizing the global variation in gene expression profiles between the control (red) and SBC8803-treated (blue) groups. The analysis was performed using normalized transcripts per million (TPM) quantification data. The first and second principal components (PC1 and PC2) explain 23% and 16% of the total variance, respectively. Each dot represents an individual biological replicate.

**Figure S3**
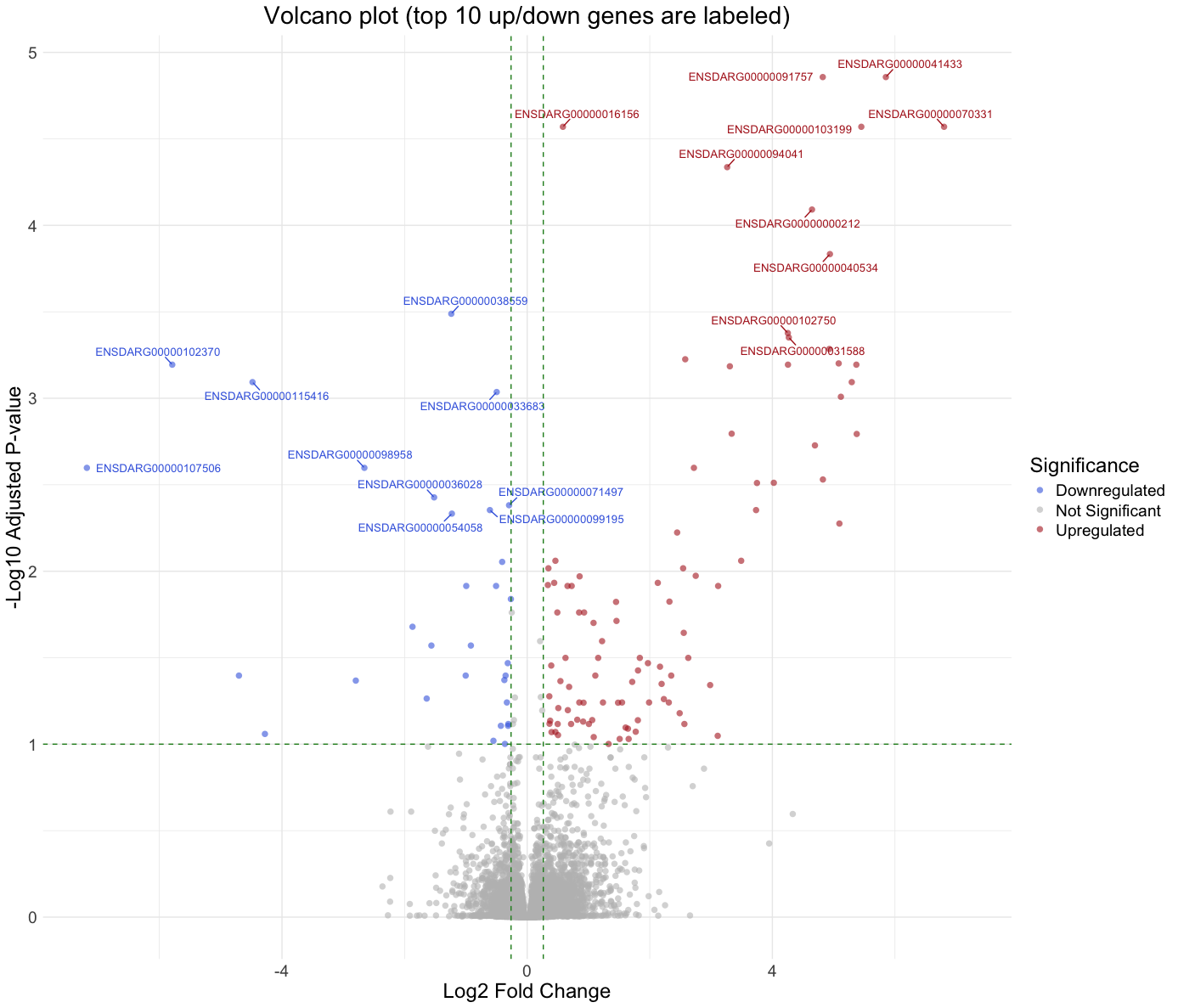

**Figure S3. Volcano plot of differentially expressed genes (DEGs) between the control and SBC8803-treated groups.** Thresholds for differential expression (adjusted *p*-value < 0.1 and |fold change| ≥ 1.2) are indicated by green dashed lines. The top 10 upregulated and downregulated genes are highlighted and labeled based on their adjusted *p*-values.

**Figure S4**

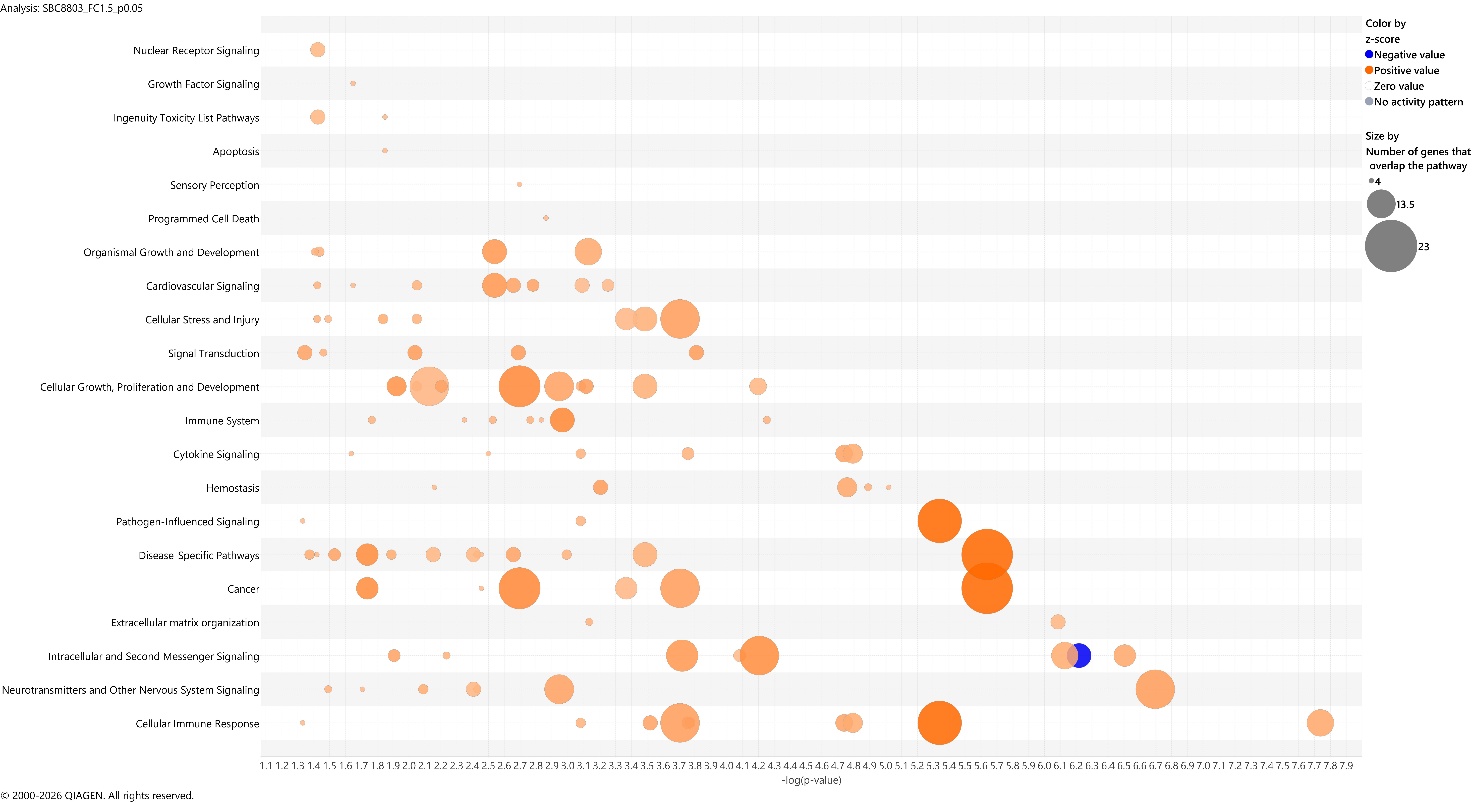

**Figure S4. Canonical signaling pathways modulated by SBC8803 administration.** Ingenuity Pathway Analysis (IPA) was performed using the differentially expressed genes to identify signaling pathways significantly enriched in the brain of SBC8803-treated zebrafish. The bar chart displays canonical pathways ranked by statistical significance. The x-axis represents the –log_10_(*p*-value) calculated by Fisher’s exact test, with a threshold of 1.3 corresponding to *p* < 0.05. Pathways with a positive z-score (predicted activation) are often depicted in orange, while those with a negative z-score (predicted inhibition) are in blue.

**Figure S5**

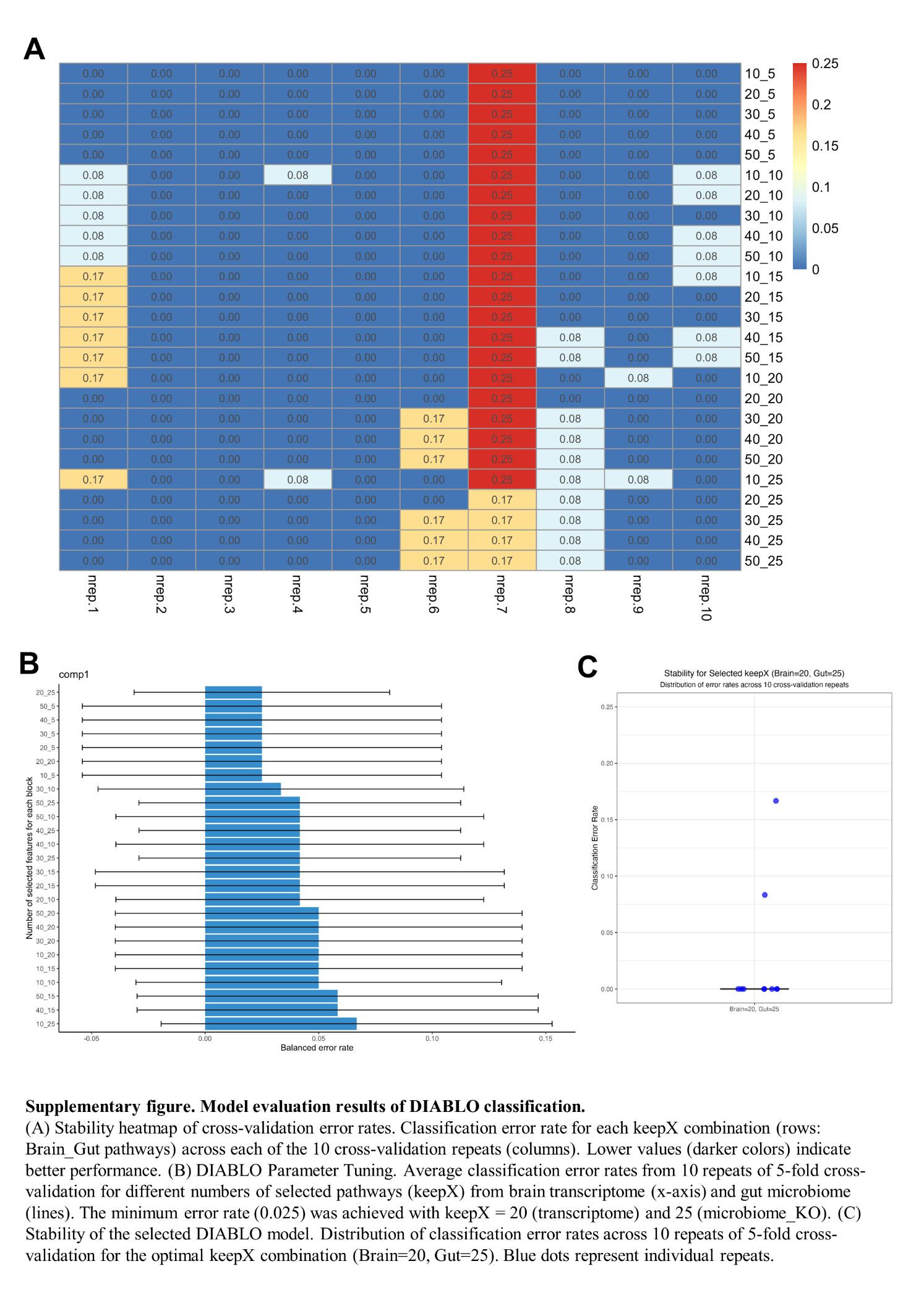

**Figure S5. Model evaluation and parameter tuning for the DIABLO classification.** **(A)** Stability heatmap of cross-validation error rates. The heatmap displays the classification error rate for various combinations of selected feature numbers (*keepX*) across 10 repeated cross-validation runs. Rows represent different *keepX* combinations for brain transcriptomic and gut microbial pathways, while columns represent individual cross-validation repeats. Lower values (darker colors) indicate better classification performance. **(B)** DIABLO parameter tuning results. The plot shows the average classification error rates obtained from 10 repeats of 5-fold cross-validation across different numbers of selected pathways (*keepX*) for the brain transcriptome (x-axis) and gut microbiome (colored lines). The minimum error rate (0.025) was achieved with an optimal architecture of *keepX* = 20 for the transcriptome and *keepX* = 25 for the microbiome. **(C)** Stability of the optimized DIABLO model. The dot plot illustrates the distribution of classification error rates across the 10 repeats of 5-fold cross-validation using the optimal *keepX* combination (Brain = 20, Gut = 25). Blue dots represent individual error rates for each repetition, demonstrating the robustness of the model.

**Figure S6**

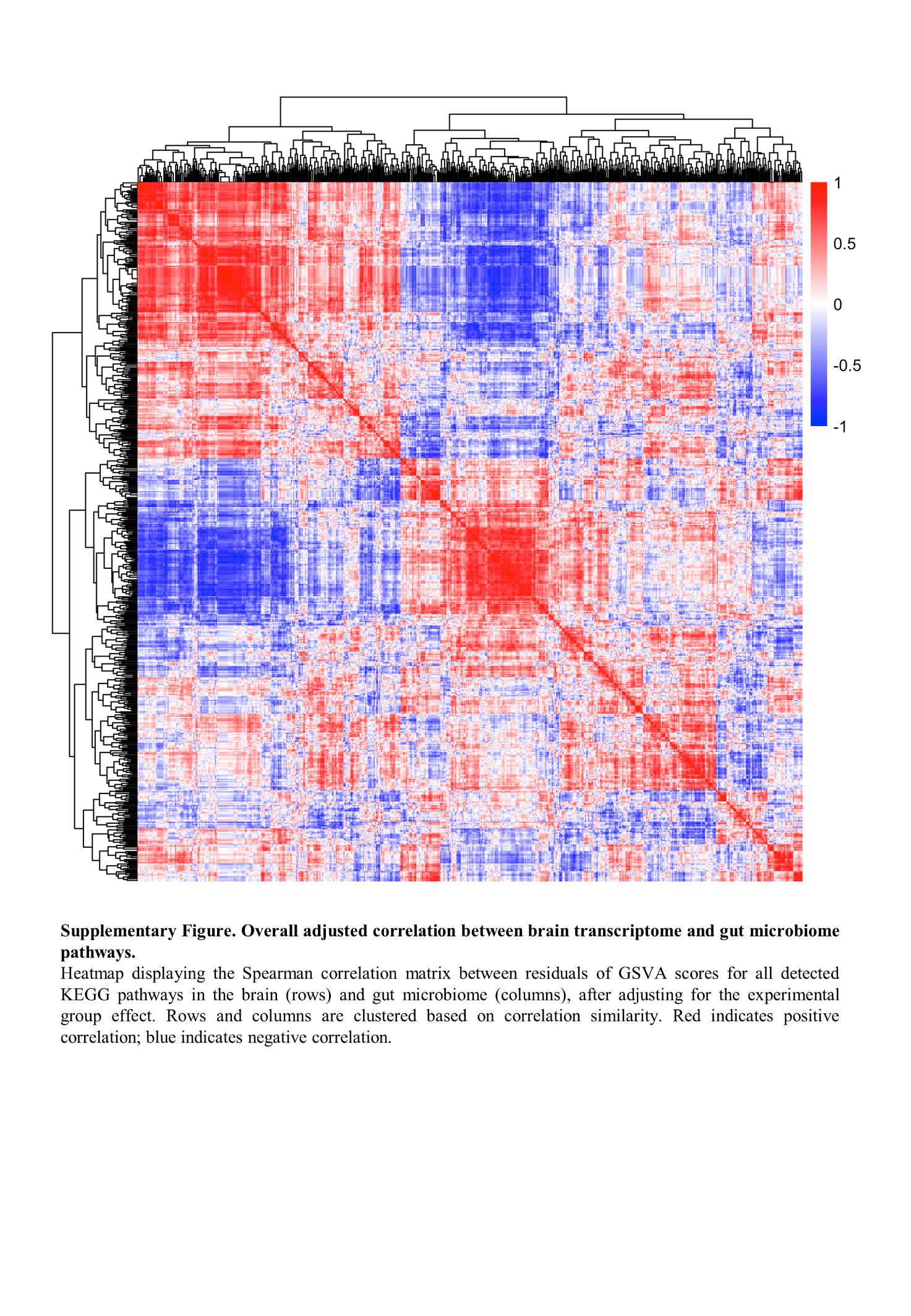

**Figure S6. Overall adjusted correlation between brain transcriptome and gut microbiome pathways.** Heatmap displaying the Spearman correlation matrix between residuals of GSVA scores for all detected KEGG pathways in the brain (rows) and gut microbiome (columns), after adjusting for the experimental group effect. Rows and columns are clustered based on correlation similarity using hierarchical clustering. Red indicates a positive correlation, while blue indicates a negative correlation. This analysis reveals the global landscape of gut–brain functional coordination independent of the dietary intervention.

**Figure S7**

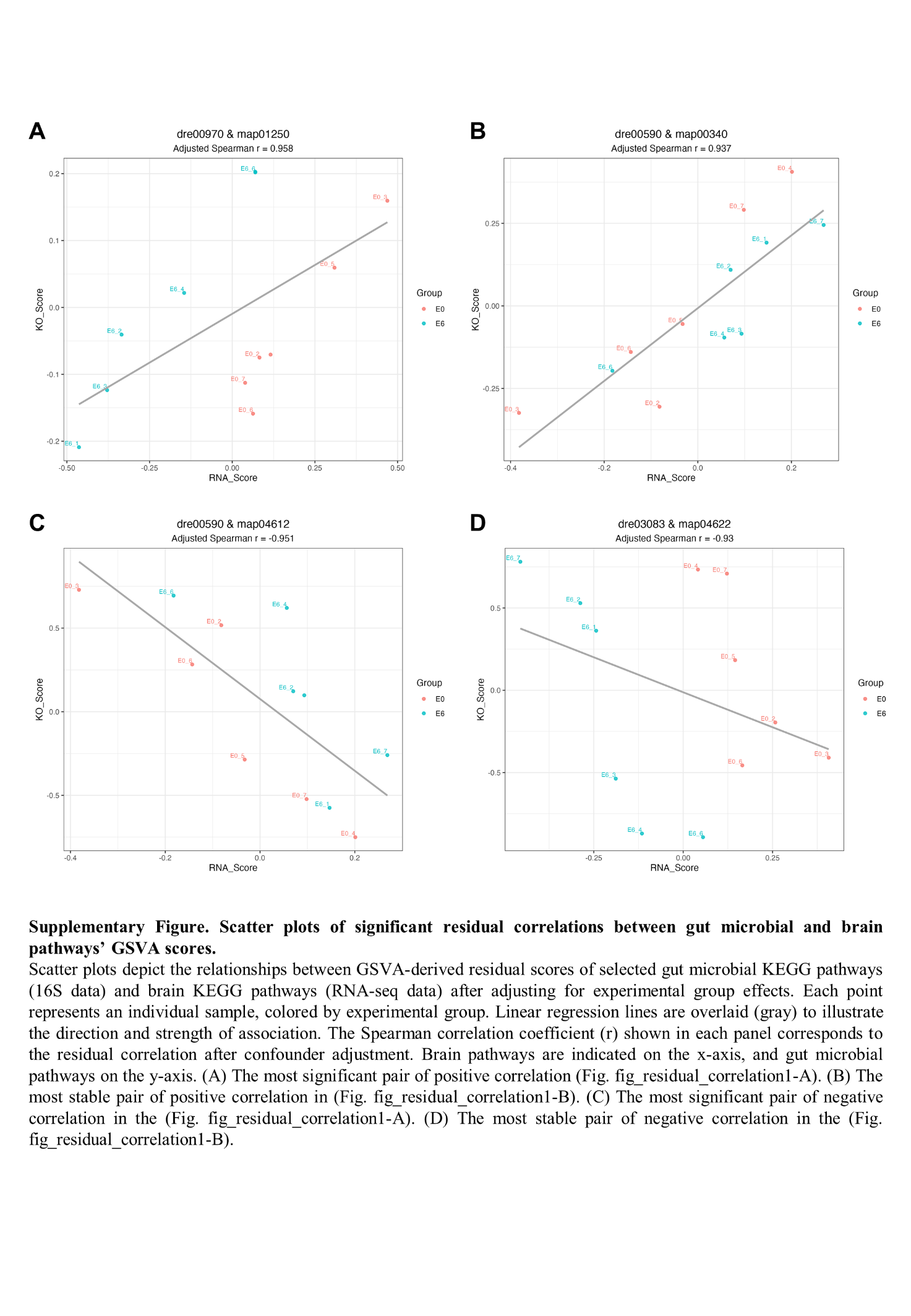

**Figure S7. Scatter plots of significant residual correlations between gut microbial and brain pathways.** Scatter plots depict the relationships between GSVA-derived residual scores of selected gut microbial KEGG pathways (y-axis) and brain KEGG pathways (x-axis) after adjusting for experimental group effects. Each point represents an individual sample. Linear regression lines are overlaid in gray to illustrate the direction and strength of the association. The adjusted Spearman correlation coefficient (*r*) is shown in each panel. **(A)** The pair with the most significant positive correlation. **(B)** The pair with the most stable positive correlation identified by sensitivity analysis. **(C)** The pair with the most significant negative correlation. **(D)** The pair with the most stable negative correlation identified by sensitivity analysis.
